# Single-cell RNA-seq reveals altered plasma cell subsets and decreased cytotoxicity of NK cells in patients with Kawasaki disease

**DOI:** 10.1101/2025.02.03.636324

**Authors:** Yuanlin Zhou, Yan Liu, Li Liu, Xiaojia Shen, Xiaobin Wen, Guixu Lin, Lin Tang, Qian Li, Jingqiu Gan, Ruixu Li, Jin Shang, Lei Li

## Abstract

**Aim:** The pathogenesis and therapeutic strategies of Kawasaki disease (KD) are worth further exploring. Using single-cell RNA sequencing (scRNA-seq), our study aimed to explore the landscape of peripheral blood mononuclear cells (PBMCs) in patients with KD.

**Methods:** We analyzed the scRNA-seq data of three patients with KD and three healthy controls from the GSE168732 dataset. Additionally, we explored the immune profiles and immune cell subtypes of patients with KD.

**Results:** Significantly increased B cells and decreased NK cells were observed in the KD group compared with that in the control group. Furthermore, six B cell subsets were identified in the PBMCs of patients with KD. Particularly, plasma cells (CD9^**+**^ and S100A9^**+**^ B cell clusters) distinctly increased in the KD group. CD9^**+**^ B cell cluster was characterized by vascular endothelial growth factor (VEGF) signaling-associated marker genes and DEGs. Meanwhile the S100A9^**+**^ B cell cluster was characterized by platelet aggregation marker genes. As for NK cells, CD16^**+**^CD56^dim^ NK cells were enriched in the KD group. Cytotoxicity-related activating receptors (KLRD1,KLRC3) and cytotoxicity-associated genes (PRF1, NKG7, GZMA, GZMH, GNLY) were downregulated in the KD group.

**Conclusion:** CD9^**+**^ and S100A9^**+**^ plasma cells may contribute to CALs in KD by upregulating VEGF and activating platelet. In addition, decreased cytotoxicity of NK cells mediated by KLRD1/KLRC3 receptor may play a vital role in the development of KD. Our results suggested that plasma cells and NK cells could be promising targets for the treatment of KD.

## 1. Background

Kawasaki disease (KD) is a multisystemic vasculitis inflammatory syndrome first described in 1967 by Rife^[1]^. KD predominantly occurs in children <5 years of age, with a reported incidence of 13.7/100000–29.8/100000^[2]^. The pathology of KD is characterized by systemic vasculitis. KD mainly affects small- and medium-sized arteries, leading to multiple organ and tissues lesions^[1, 3]^. Approximately 30–50% of patients with KD have complicated coronary artery dilation, and approximately 25% have serious coronary artery damage, including coronary artery aneurysm and ectasia. Artery dilatation or aneurysm caused by KD is considered as the most common acquired cardiovascular disease in children^[2, 4]^. Coronary artery damage directly affects the severity and long-term prognosis of KD. Severe coronary artery damage leads to a worse long- term prognosis, myocardial infarction, and sudden death. Most studies have characterized KD as an abnormal activation of the immune system in genetically susceptible children, which is triggered by infectious factors^[1, 5, 6]^. In the pathogenesis of KD, dysregulated innate and adaptive immune responses lead to arterial infiltration by neutrophils, CD8^**+**^ T cells, B cells, dendritic cells, and monocytes/macrophages. Furthermore, inflammatory signaling cascades are activated. The NF-κB signaling pathway is the classical pro-inflammatory signaling mechanism in KD. Binding of ligands to receptors (such as tumor necrosis factor, pattern recognition, and interleukin-1) can activate the NF-κB signaling pathway in KD, resulting in translocation of the activated NF-κB to the nucleus and triggering of abundant proinflammatory cytokines and chemokine expression, including that of NOD-like receptor family pyrin domain containing 3 (NLRP3); interleukin (IL)-1, - 6, -8, -17, and 18; tumor necrosis factor α (TNF-α); interferon γ (IFN-γ); and vascular endothelial growth factor (VEGF)^[7-11]^. Additionally, NF-κB activation had been reported to promote the secretion of inflammatory cytokines by macrophages, following which, vascular inflammation was induced by an imbalance in Th1/Th17 polarization and recruitment of myeloid cells. Consequently, vascular endothelial cell injury eventually develops. Moreover, several studies have reported that the activation of the VEGF signaling pathway plays a crucial role in the development of coronary artery lesions (CAL) in KD. Evidence indicates that VEGF levels strongly correlate with CAL severity in KD with VEGF acting directly on endothelial cells, resulting in increased blood vessel permeability and fibrin deposition. This pathological process leads to extracellular matrix degradation and endothelial dysfunction in patients with KD.

Currently, intravenous immunoglobulin gamma (IVIG) combined with oral aspirin is the predominant treatment for KD. IVIG treats KD through several immunomodulatory mechanisms. On one hand, IVIG inhibits the activation of the innate immune system by interacting with various cell types, including neutrophils; macrophages; and NK, dendritic, and vascular endothelial cells. On the other hand, IVIG suppresses the activation of adaptive immunity by interacting with T cells and antibodies. The antigen-binding fragment (Fab) of IVIG suppresses Th17 differentiation and proliferation, and pro-inflammatory cytokine expression. The heavy constant region (Fc) of IgG recognizes Tregs to regulate adaptive immunity in KD. In addition, IVIG can interact with molecules such as PAMPS/DAMPs, NLRP3, NLRP1, and IkBα to suppress the activation of the innate immune system in KD. However, approximately 10–18% of patients with KD are resistant to IVIG therapy. Nonresponders to IVIG have a high risk of developing coronary artery aneurysms. The incidence of coronary artery aneurysms in IVIG nonresponders is reported to be as high as 31.3%^[12]^. Although methylprednisolone, the TNF-α monoclonal antibody infliximab, cyclophosphamide, methotrexate, cyclosporine A, plasmapheresis, and plasma exchange are supplementary therapies for IVIG-resistant KD^[11, 13]^, approximately 4.6% patients experience coronary artery aneurysms after therapy^[11, 14]^. Hence, the specific mechanism of KD is worth exploring to improve the efficacy of IVIG.

Recently, single-cell RNA sequencing (scRNA-seq) has been used to reveal cell heterogeneities in many immune diseases. The mechanisms resulting in aberrant immune function or overexpression of inflammatory cytokines in KD need to be explored. In this study, we employed scRNA-seq to analyze lymphocyte subtypes and reveal the heterogeneities of different lymphocyte subtypes in KD, with the aim of providing new insights into the mechanism of KD and its treatment targets.

## 2. Materials and Methods

### 2.1 Data sources

The scRNA-seq files (GSE168732 dataset) for 3 KD samples and 3 healthy control samples were download from the GEO database (https://www.ncbi.nlm.nih.gov/geo/). 3 KD samples all were pre-treatment samples.

### 2.2 Preprocessing of single-cell RNA-seq data and quality control

Primitive sequencing data of GSE168732 dataset were processed using the Cell Ranger pipeline(version 5.0.0), including cell barcode and unique molecular identifier (UMI) counting. The UMI was counted using Seurat20 for quality control. The criteria applied for quality control were as follows: total UMI count > 1000, log10 GenesPerUMI > 0.7, mitochondrial gene percentage < 5% and hemoglobin gene percentage < 5%. Doublet droplet were eliminated by Doublet Finder package (version 2.0.2).Mutual nearest neighbors (MNN) was used to reduce the dimensionality. Cell community and clustered cells were established through Shared Nearest Neighbor (SNN). Uniform manifold approximation and projection (UMAP) was performed to visualize cell clusters.

### 2.3 Marker genes and cell subsets identifying

Marker genes of each clusters were identified through FindAllMarkers function. R package SingleR was used to recognize the cell types of scRNA-seq.

### 2.4 Identification of DEGs and functional analysis

Differentially expressed genes (DEGs) between groups of different cell types were identified using the Seurat Find Marker function with the default parameters. The criteria for identifying DEGs was an average log2 fold change > 0.25. The cutoff adjusted *p*-value was set at 0.05. Gene ontology (GO) and Kyoto Encyclopedia of Genes and Genomes (KEGG) were used to analyze biological processes, pathways, and functions of the DEGs.

### 2.5 Ethics approval and consent to participate

The study was approved by the Ethics Committee of Chengdu Second People’s Hospital. This study follows the Declaration of Zhen Wang^[15]^.

## 3. Results

### 3.1 Composition of different immune cells and DEGs in KD and healthy children

The scRNA-seq data of three patients with KD and three healthy children from the GSE168732 dataset were analyzed to reveal the immune features of KD. In total, 37090 cells were detected in six samples after quality control. Ten major celltypes were identified in the PBMCs according to the canonical marker gene using UMAP: B cells, macrophage-monocytes, plateles, naïve CD4^**+**^ T cells, naïve CD8^**+**^ T cells, Th1 cells, effector T cells, active T cells, NK cells, and neutrophils (Figures 1A and 1B). Moreover, sixteen PBMCs clusters were identified in the KD and control groups, including Tfh, Th1, Tc17, active T, plasma, and naïve B cells; classical monocytes; M1 macrophages; platelets; CD56^dim^ NK cells; and neutrophils.

**Figure 1.**
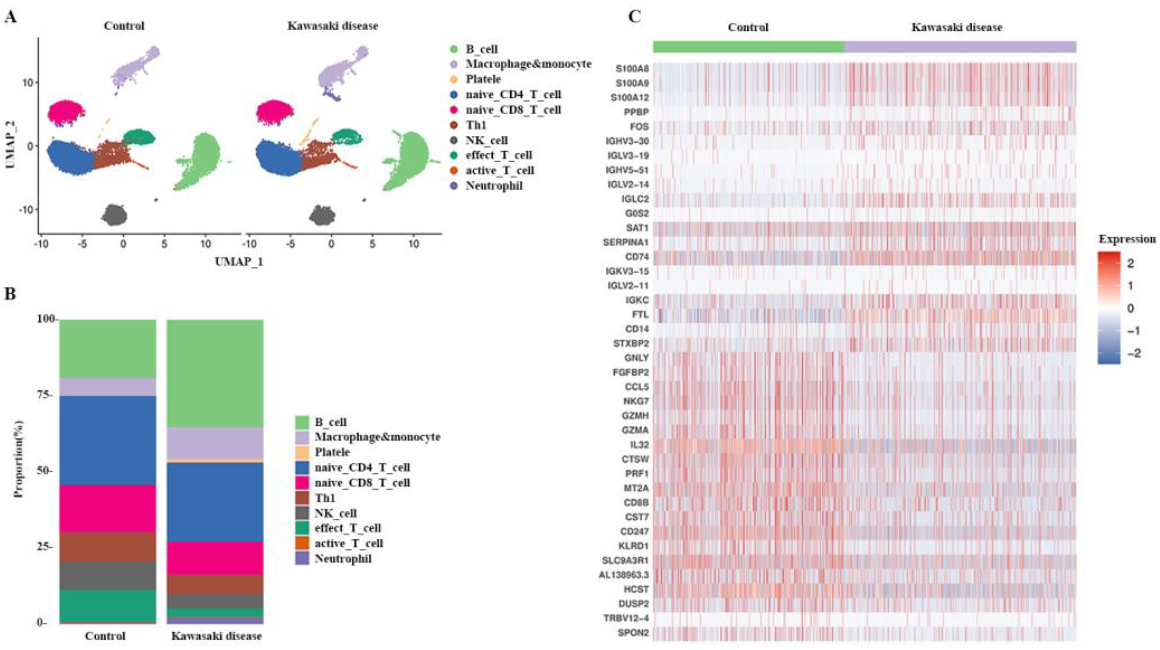
Clustering of peripheral blood mononuclear cells (PBMCs) in Kawasaki disease (KD). (A) Ten major celltypes were identified in PBMCs; (B) Differences in the composition of 10 major PBMC celltypes between the KD and healthy control groups; (C) Heatmap of top 40 differentially expressed genes (DEGs) identified in PBMCs between the KD and healthy control groups

We also analyzed PBMC distribution compositions, which were generally different between the KD and health control groups (Figure 1B). An increasing trend was observed in platelets, B cells, monocytes, and neutrophils while a decreasing trend in naive CD8^**+**^ T, effector T, and NK cells in the KD group compared with the control group. Furthermore, DEGs among all annotated cellular types were identified in children with KD and in healthy children. A total of 94 DEGs were detected between the two groups (*p* < 0.05, FoldChange > 1.5). Among these, the top 40 DEGs included S100, immunoglobulin, and granzyme family coding genes (Figures 1C). We combined the results of DEGs with those of GO and KEGG pathway enrichment analyses(Figure 2). The results showed that three of the top five significantly upregulated genes: S100A8, S100A9, and FOS, participated in the IL-17 signaling pathway, which plays an important role in Th17 differentiation in KD. Moreover, five significantly upregulated genes: IGHV3-30, IGHV5-51, IGKC, and IGLC2, participated in the immunoglobulin-mediated immune response, indicating abnormal secretion of immunoglobulin. In contrast, : CD247, HCST, KLRD1, and PRF1, normally participate in NK cell-mediated cytotoxicity. This indicated a dysfunction of NK cells in KD. Notably, we found that the ferroptosis regulatory genes FTL and SAT1 were both upregulated in KD.

**Figure 2.**
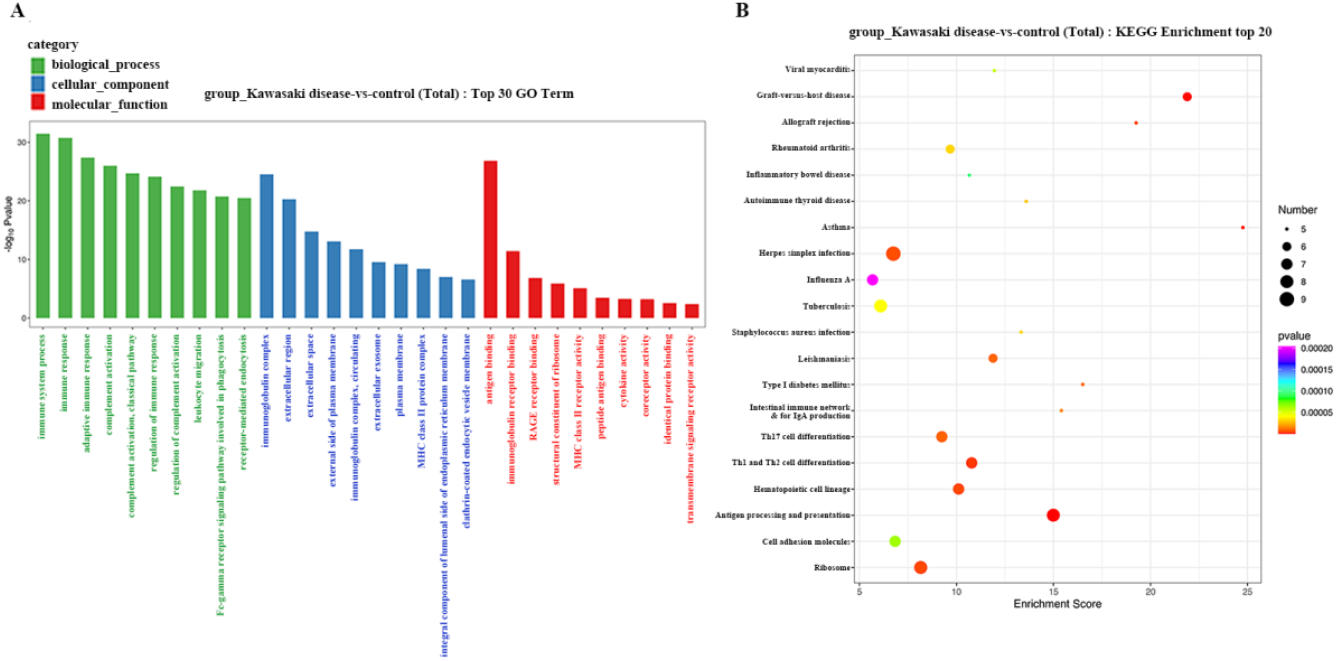
Association of peripheral blood mononuclear cells (PBMCs) with Kawasaki disease (KD). (A) Top 30 Gene Ontology (GO) terms in differentially expressed genes (DEGs) identified in PBMCs between the KD and healthy control groups; (B) Top 20 enrichment pathways according to Kyoto Encyclopedia of Genes and Genomes (KEGG) analysis of DEGS identified in PBMCs between the KD and healthy control groups.

These results indicate that an increase in B cells, B cell-mediated platelet activation, IL-17 signaling pathway activation, NK cell-mediated cytotoxicity inhibition, and VEGF-mediated angiogenesis play important roles in the development of KD.

### 3.2 Association between different cell types and KD development

We further explored the heterogeneity and potential functions of different cell types in KD development. Specific marker genes in different cell types were identified using the FindAllMarkers function and DEGs were analyzed using KEGG and GO analyses.

### 3.2.1 CD9^**+**^ and S100A9^**+**^ B cell clusters are associated with development of CAL in KD

The increasing trend in the number of B cells was particularly significant. We further subdivided the B cell subsets according to canonical marker genes (Figure 3). Six B cell subsets were identified (Figure 3A). The B cell subsets distribution displayed a distinct pattern in patients with KD and in healthy individuals (Figure 2B). Compared with the control group, patients with KD exhibited an increasing trend in the CD9^**+**^ B cell (cluster 3 in Figure 3A) and S100A9^**+**^ B cell (cluster 6 in Figure 3A) clusters. Patients with KD showed decreasing trends in the naïve (cluster 1 in Figure 3A) and memory B cell (cluster 4 in Figure 3A) clusters. The CD9^**+**^ and S100A9^**+**^ B cell clusters are two novel B cell subsets enriched in plasma cells. The CD9^**+**^ B cell cluster was characterized by the expression of angiogenesis-associated marker genes (CD9, PIK3IP1, PELI1,KLF2, HERPUD and EZR).Meanwhile, angiogenesis-associated marker genes KLF2, HERPUD1, EZR, and CD9 were also the significant DEGs in B cells between KD group and control group(supplementary in Figure 4). In addition, S100A9^**+**^ B cell cluster was characterized by marker genes involved in platelet aggregation (S100A8,S100A9,GP9, TGA2B, ITGB1, TREML1, PPBP, and PF4). S100A8 and S100A9 were distinctly increased in KD group (supplementary in Figure 4). The S100A9^**+**^ B cell cluster is a novel B cell subset identified in patients with KD.

**Figure 3.**
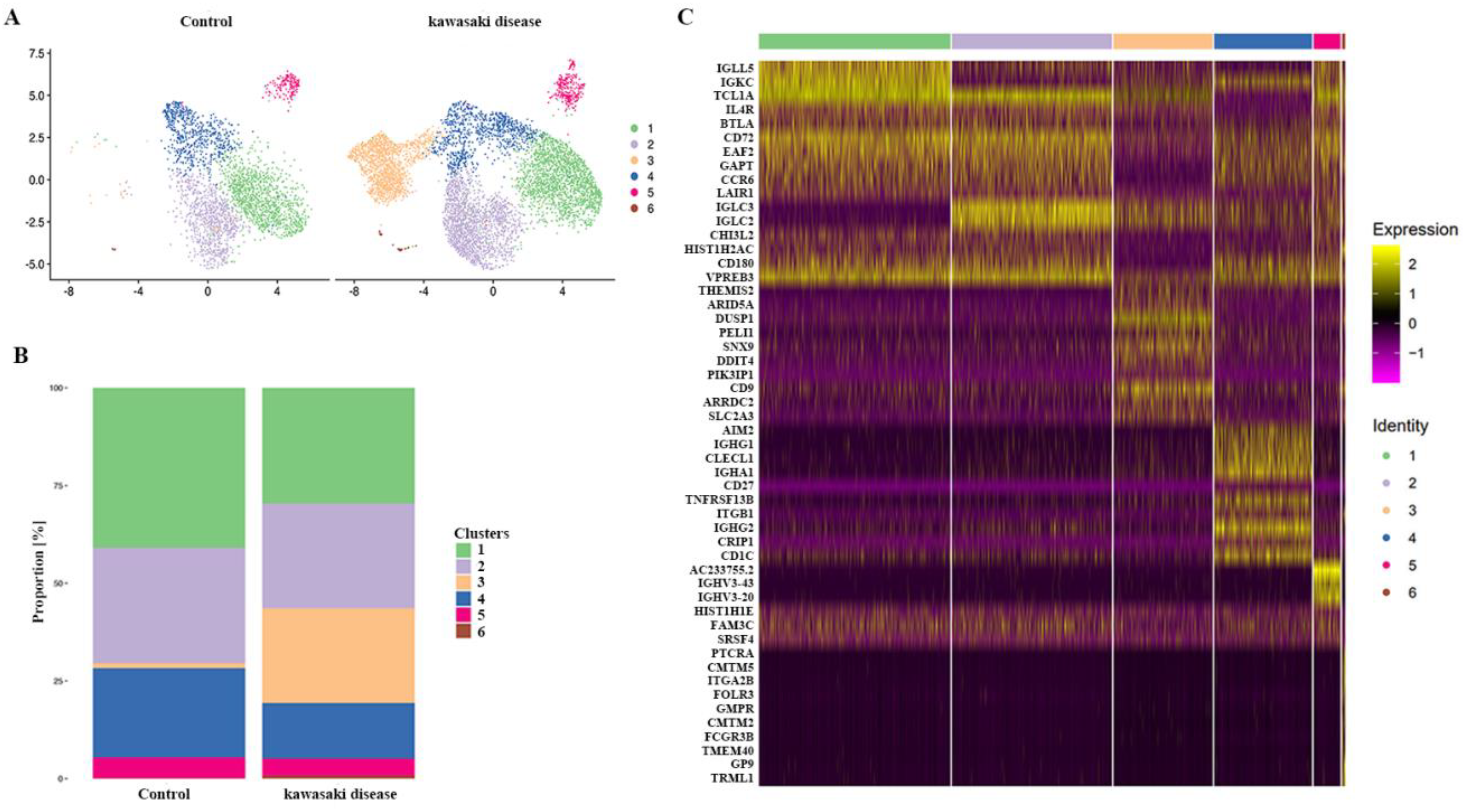
Association of B cells with Kawasaki disease (KD) development. (A) Uniform manifold approximation and projection (UMAP) plot of six B cell clusters; (B) Differences in the composition of six B cell clusters between the KD and control healthy groups. (C) Heatmap of unique signature genes in each B cell cluster.

Regarding the DEGs in B cells, seven of the top 16 significant DEGs were involved in vascular endothelial dysfunction and angiogenesis, including EZR, CD9, HERPUD1, MEF2C, and KLF2. The DEGs upregulating VEGF and accelerating angiogenesis (EZR, HERPUD1, and CD9) were upregulated in the KD group. The angiogenesis suppression DEGs MEF2C and KLF2 were downregulated in the KD group. Furthermore, the immunoglobulin-associated DEGs GLC2 and IGLC3,which identified as marker genes in CD9^**+**^ B cell cluster, were significantly upregulated in KD group (Figure 4).

**Figure 4.**
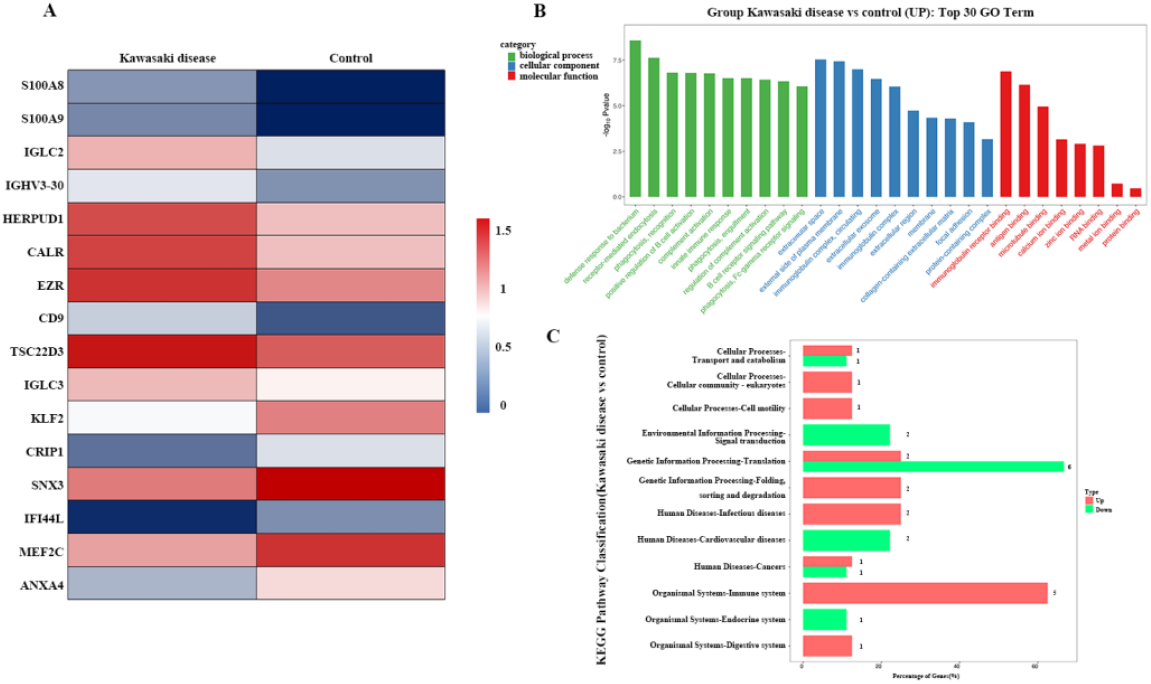
Association of B cells with Kawasaki disease (KD). (A) Heatmap of the significant differentially expressed genes (DEGs) identified in B cells between the KD and healthy control groups; (B) Top 30 Gene Ontology (GO) terms of DEGs identified in B cells between the KD and healthy control groups; (C) Top 20 enrichment pathways according to Kyoto Encyclopedia of Genes and Genomes (KEGG) analysis of DEGs identified in B cells between the KD and healthy control group

#### 3.2.2 Decreased cytotoxicity of NK cells in KD

Cell subtye analysis indicated that CD16^**+**^CD56^dim^ NK cells were the major NK cells enriched in patients with KD. CD16^**+**^CD56^dim^ NK cells are cytotoxic. CD16^**+**^CD56^dim^ NK cells identified in our study expressed non-HLA-specific activating receptors (CD16, KLRF1, KLRD1/KLRC3, and NCR1), HLA-class-⍰ activating receptors KLRD1/KLRC3 and the HLA-class-⍰ NK cell inhibitor receptor (KLRD1/KLRC1, KIR2DL3, and KIR3DL2) (also named CD94/NKG2A). The binding of NK cell activating receptors to their ligands promotes the killing ability of NK cells and increases the secretion of inflammatory cytokines such as IFN-γ and TNF-α. The binding of inhibitor receptors (CD49/NKG2A, KIR2DL3, and KIR3DL2) to their corresponding ligands inhibits NK cell function^[16]^. KLRD1 forms a heterodimeric with activating receptors NKG2E and activats the cytotoxicity of NK cells. KLRD1 was signifcantly decreased in PBMCs of KD group while NKG2E was boviously downregulated in NK cell clusters of KD group (Figure 1C and Figure 5A). In addition, CD16^**+**^CD56^dim^ NK cells enriched in patients with KD were characterized by the cytotoxicity-associated marker genes and dereased DEGs GZMA, GZMH, GNLY, FGFBP2, PRF1, and NKG7 (Figure 5A). In addition, the downregulated DEG MT2A is involved in protecting the heart from cardiotoxic drugs. NK cell proliferation and survival-related genes, such as KLF2, ZNF683, and MALAT1, were also downregulated. These results indicate that inhibition of cytotoxicity occur in NK cells during KD development (Figure 5A).

**Figure 5.**
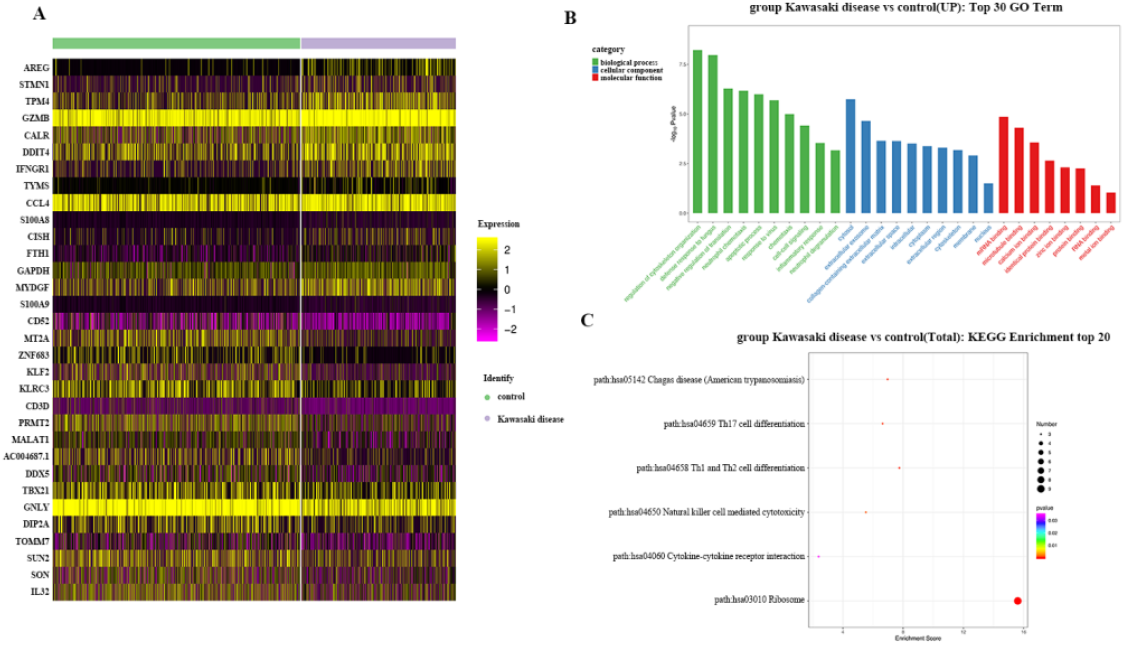
Association of NK cells with Kawasaki disease (KD). (A) Heatmap of the significant differentially expressed genes (DEGs) identified in NK cells between the KD and healthy control groups; (B) Top 30 Gene Ontology (GO) terms enriched in DEGs identified in NK cells between the KD and healthy control groups; (C) Heatmap of the top 20 pathways enriched by DEGs identified in NK cells between the KD and healthy control groups

Moreover we found that five of the 15 most significantly upregulated DEGs, including AREG, S100A8, S100A9, CCL4, and TYMS, were involved in the positive regulation of the inflammatory response. IFNGR1 is also involved in IFN-γ signaling. These genes are related to the pro-inflammatory effects of NK cells. Moreover, the upregulated DEGs TPM4, CALR, and STMN1 were associated with vascular diseases. STMN1 is involved in ferroptosis in coronary artery disease. TPM4 participates in vasculopathy-related striated muscle contraction, whereas CALR participates in calcium regulation in cardiac cells. DDIT4 is involved in autophagy. These genes are related to the abnormal programmed death of cardiac cells (Figure 5A).

### 3.3 CD177^**+**^ neutrophils are associated with KD

Many studies have demonstrated that neutrophil infiltration of the coronary artery wall is a classic innate immune response associated with the development of KD. In our study, we found that the proportion of neutrophils was markedly increased in the KD group. To explore the function of neutrophils in KD, marker genes and DEGs in neutrophils were identified between the KD and healthy groups (Figure 6). CD177^**+**^ neutrophils were enriched in patients with KD. CD177, which promotes the adhesion of neutrophils to the vascular wall through interaction with PECAM-1 and β2 integrins, were significantly higher in the KD group than in the control group. Two of the top 10 marker genes (ADGRG3 and CD177) are components of the PR3/CD177/GPR97/PAR2/CD16b complex.. FCGR3B was identified as a significant gene marker and was noticeably increased in neutrophils in the KD group compared with the control group. Moreover, many receptors were identified as significant marker genes and were distinctly upregulated in the neutrophils cluster of the KD group, including the FLMP (FPR1, SOD2), CXCL chemokine (CXCL8, CXCL1, NAP-2) and opsonin (FCGR2A, FCGR3A, FCGR3B, ITGAM, CR1) receptors. In addition, three of the top 10 marker genes associated with matrix deposition in vasculitis (ALPL, MMP25, ANXA3) were increased in the KD group. Notably,eight of the top 40 significant DEGs associated with vasculitis: CXCL8, PROK2, SAT1, ANXA3, EGR1, ADM, S100A8, and S100A9, were upregulated in the KD group. Furthermore, IFN- and M1 macrophage-stimulation genes, including EGR1, IFITM2, IFITM3, SAT1, and NEAT1, were enriched in the neutrophil cell cluster of the KD group. 12 of the top 40 significantly upregulated DEGs (S100A8, S100A9, S100A11, S100A12, CD177, FCGR3B, CXCL8, FOLR3, FPR1, PGLYRP1, PLAUR, and PROK2) were involved in neutrophil degranulation and cytotoxicity.

**Figure 6.**
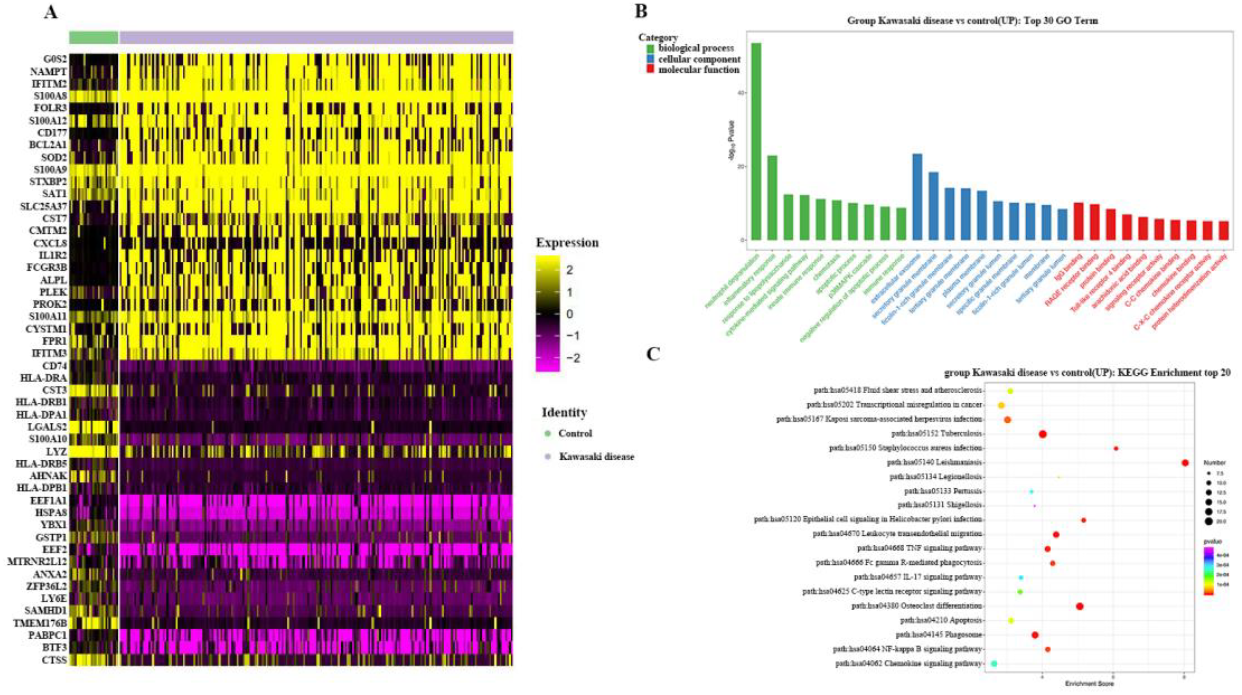
Association of neutrophils with Kawasaki disease (KD). (A) Heatmap of the top 40 significant differentially expressed genes (DEGs) identified in neutrophils between the KD and healthy control groups; (B) Top 30 Gene Ontology (GO) terms of DEGs identified in neutrophils between the KD and healthy control groups; (C) Heatmap of the top 20 enrichment pathways in DEGs identified in neutrophils between the KD and healthy control groups

### 3.4 Correlation of monocytes/macrophagocyte and T cells with KD

We also explored the phenotypic heterogeneity of monocytes/macrophages and T cells in patients with KD. Three subsets of macrophages (macrophage_VCAN, macrophage_C1QA, and macrophage_s100A12) and five subsets of T cells (Tfh, Th1, naive Tc17, Tc17, and active T cells) were identified in the PBMCs. Three subsets of macrophages were characterized by HLA class II neoantigens and CD68 marker genes, indicating that pro-inflammatory M1 macrophages release many pro-inflammatory cytokines and promote the differentiation of Th1 and Th17 cells in KD. Regarding DEGs in monocytes/macrophages in KD, the five most significantly upregulated and marker genes in monocyte/macrophagocyte clusters: S100A8, S100A9, S100A12, RETN, and CLU, noticeably participated in NF-κB activation and positively regulated inflammatory responses. Among the upregulated DEGs, FOS mainly participated in the activation of the IL-17 signaling pathway; NCF4 is involved in phagosomes; and PIM1 is involved in the JAK-STAT signaling pathway, which indicates that monocytes/macrophages in KD exhibit pro-inflammatory features. In addition, the coronary artery disease-related genes MCEMP1, VCAN, TXNIP, and CD74 were identified as marker genes and DEGs in the monocyte/macrophagocyte cluster (Figure 7).

**Figure 7.**
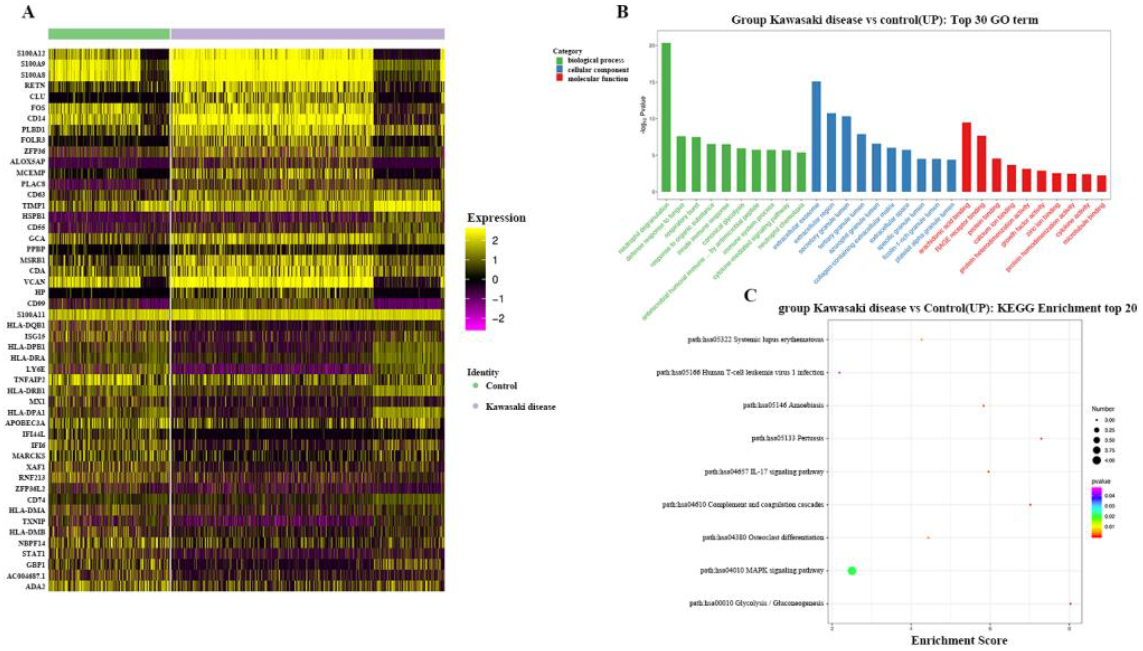
Association of monocytes/macrophagocyte cells with Kawasaki disease (KD). (A) Heatmap of the significant differentially expressed genes (DEGs) identified in monocytes/macrophagocyte cells between the KD and healthy control groups; (B) Top 30 Gene Ontology (GO) terms of DEGs identified in monocytes/macrophagocyte cells between the KD and healthy control group; (C) Heatmap of the top 20 enrichment pathways of DEGs identified in monocytes/macrophagocyte cells between the KD and healthy control group

Among the top 25 DEGs of T cells in KD, the four most significantly upregulated genes: CISH, PIM1, MYC, and SOCS3, were all involved in the regulation of the JAK-STAT signaling pathway, which is associated with macrophage polarization. In contrast, cytotoxicity-related genes, including GNLY, GZMH, NKG7, FCGR3A, GZMA, and KLRD1, were significantly downregulated in the T cells of patients with KD.

## 4. Discussion

Kawasaki disease has been reported to be associated with overactivation of the immune response and inflammatory storms. However, its pathology remains unclear. To explore the immune response in KD, we used scRNA-seq to comprehensively describe the immune landscape of PBMCs from patients with KD. Furthermore, we analyzed the composition and effects of immune cells in patients with KD.

In this study, we found that B cells are associated with angiogenesis and platelet activation in KD development. The CD9^**+**^ B cell cluster expressed the VEGF-associated marker genes D9, PIK3IP1, PELI1,KLF2, HERPUD and EZR. VEGF-associated DEGs were identified in KD group and control group. There is mounting evidence that the upregulated expression of VEGF, VCAM-1, and ICAM-1 promotes CAL in KD and plays a crucial role in the mechanisms of KD^[17, 18]^, suggesting that B cells may promote CAL of KD by activating VEGF signaling pathway. In addition, we identified a novel B cell subset (S100A9^**+**^ B cell cluster) in patients with KD. The S100A9^**+**^ B cell cluster was characterized by marker genes involved in platelet aggregation, including GP9, TGA2B, ITGB1, TREML1, PPBP, and PF4. This phenomenon indicates that platelets may interact with platelet receptors on S100A9^**+**^ B cell clusters. Furthermore, the S100 family proteins S100A8 and S100A9 were identified in the S100A9^**+**^ B cell cluster and were significantly upregulated in the KD group. S100A8 and S100A9 are also associated with KD and other cardiovascular diseases such as atherosclerosis and myocardial infarction^[19, 20]^. In contrast, S100A8 and S100A9 drive the formation of pro-coagulant platelets^[21-24]^. Moreover, S100A8 and S100A9 can induce the expression of ICAM-1 and VCAM-1 on endothelial cells^[25, 26]^. S100A8 and S100A9 can also activate the NF-κB signaling pathway[25, 27, 28]. This result suggests that the CD9^**+**^ B cell cluster may upregulate VEGF expression, whereas the S100A9^**+**^ B cell cluster may interact with platelets in KD. We identified distinct B cell subset features between patients with KD and healthy children. In addition, our results reveal the underlying role of B cells in the development of CAL in patients with KD.

Furthermore, we found that CD56^dim^ NK cells with activating receptors (CD16, KLRF1, KLRD3/KLRC3, and NCR1) and HLA-specific NK cell inhibitor receptors (KLRD1/KLRC1, KIR2DL3, and KIR3DL2) were enriched in patients with KD^[16, 29]^. NK activating receptors genes KLRD1/KLRC3 and cytotoxicity-associated genes (PRF1, GZMA, GZMH, GNLY,,NKG7 and FGFBP2) were downregulated in the KD group. The decreased cytotoxicity of NK cells has been demonstrated in many autoimmune diseases, such as multiple sclerosis, systemic lupus erythematosus, rheumatoid arthritis, and type-1 diabetes^[30-33]^. Decreased NK cell cytotoxicity is considered influential in the initiation and onset of multiple sclerosis and diabetes[32, 34, 35]. Insufficient NK cell cytotoxicity is reportedly associated with multiple sclerosis progression^[30, 36]^. Similarly, the proportion of CD56^dim^ NK cells showed an increasing trend, whereas cytotoxicity-associated genes (KLRD1, KLRC3, NKG7, PRF1, GZMA, GZMH, GNLY, and FGFBP2) were downregulated in the KD group. This indicates patients with KD exhibited decreased cytotoxicity of NK cells, which may be mediated by the KLRD1/KLRC3 receptors. Decreased cytotoxic activity of NK cells can result in a weakened defense against pathogenic infections. This appears to be vital in the pathogenesis of KD. This suggests that suppression of NK cell cytotoxicity contributes to KD development.

Regarding neutrophils, monocytes/macrophages, and T cells, we found an increasing trend in neutrophils and monocytes/monocytes in the KD group and a decreasing trend in naive CD8^**+**^ T, Th1, and effector T cells in the KD group. CD177^**+**^ neutrophils were predominantly enriched in the KD group and CD177 was distinctly upregulated in the KD group. These results are consistent with those of previous studies^[37, 38]^. In addition, we found that the top marker genes (GPR97 and CD177) in the neutrophil cluster were markedly increased in the KD group. GPR97 and CD177 are important components of the PR3/CD177/GPR97/PAR2/CD16b complex^[39]^. This result indicates that neutrophil activation and pro-inflammatory cytokine release may be mediated by the PR3/CD177/GPR97/PAR2/CD16b complex. However, this phenomenon has not been reported previously. Moreover, we identified monocytes/macrophages in KD characterized by HLA class II neoantigens and CD68 marker genes, suggesting that pro-inflammatory M1 macrophages release many pro-inflammatory cytokines and promote the differentiation of Th1 and Th17 cells in KD. The five most significantly upregulated genes were pro-inflammatory genes S100A12, S100A8, S100A9, RETN, and CLU. Coronary artery disease-related genes MCEMP1, VCAN, TXNIP, and CD74 were identified as marker genes and DEGs in monocyte/macrophagocyte clusters, indicating that MCEMP1, VCAN, TXNIP, and CD74 may be involved in the CAL of KD^[40-45]^. In the T cell cluster, cytotoxic genes, including GNLY, GZMH, NKG7, GZMA, KLRD1, and FCGR3A, were significantly reduced in T cells of patients with KD. This result suggests that the cytotoxicity of T cells decreased in the KD group, and resistance to pathogens was reduced during the development of KD.

## 5. Conclusion

Previous studies have focused on the pathogenesis of KD in T cells and monocytes/monocytes. Information on B and NK cells in KD is lacking. In the present study, we found that the CD9^**+**^ B cell cluster was associated with the development of CAL in KD through activation of VEGF signaling. S100A9^**+**^ B cell cluster may interact with platelets and involve in platelet aggregation during CAL pathogenesis in KD. In addition, our study revealed that CD16^**+**^CD56^dim^ NK cells are enriched in patients with KD. Decreased NK cell cytotoxicity mediated by KLRD1/KLRC3 may attenuate defense against pathogen infections and contribute to the development of KD. Our study provides a single-cell landscape of immune features and novel insights into the pathogenesis of KD, providing helpful evidence to understand the mechanism and therapeutic strategy for KD.Targeting plasma cell and increasing cytotoxicity of NK cells may be promoting options for the treatment of KD.

## List of abbreviations

KD: Kawasaki disease
PBMCs: peripheral blood mononuclear cells
scRNA-seq: single-cell RNA sequencing
VEGF: vascular endothelial growth factor
NK cell: natural killer cell
IVIG: intravenous immunoglobulin
DEGs: differentially expressed genes
scRNA-seq: single cell RNA sequencing
PBMCs: peripheral blood mononuclear cells
IL: interleukin
TNF-α: tumor necrosis factor α
UMI: unique molecular identifier
DEGs: Differentially expressed genes

## Acknowledgements

We sincerely appreciate the funding supporters of the study. In addition, we thank Chengdu Baimeisen Biological Technology Co. LTD (Sichuan, China) for scRNA bioinformatics support.

